# Progression of visual-stimulus-evoked blood velocity response timing across the vascular network measured with individual-vessel phase-contrast fMRA

**DOI:** 10.64898/2026.10.16.759420

**Authors:** Zhangxuan Hu, Grant A. Hartung, Sebastien Proulx, Divya Varadarajan, Saskia Bollmann, Bruce R. Rosen, Jonathan R. Polimeni

**Author notes:** Correspondence to: Zhangxuan Hu, PhD. Current address: Richard M. Lucas Center for Imaging, Department of Radiology, Stanford University School of Medicine, 1201 Welch Road, Stanford, CA, USA. Current address: Institute for Mechanics, Computational Mechanics Group, Technical University of Darmstadt, Darmstadt, Germany (Grant A. Hartung).

## Abstract

Most functional MRI (fMRI) studies of brain function aim to measure hemodynamic responses arising from the microvasculature within the tissue parenchyma, where spatial and temporal coupling with neuronal activity is thought to be tightest. However, the measured hemodynamic responses emerge from an interconnected vascular network that includes upstream feeding arteries, downstream draining veins, and the intervening microvascular compartments. Directly measuring and comparing vascular responses across multiple stages of this network may therefore provide more specific information about network-level hemodynamics—and how hemodynamics in one stage of the vascular hierarchy influence hemodynamics in others—that is not accessible from conventional fMRI signals alone. This can provide new insights into the relationship between the observed fMRI responses and the underlying neuronal activity of interest.

This study extends our recently proposed dynamic phase-contrast functional MR angiography (PC-fMRA) method for tracking stimulus-evoked blood-velocity responses in large supplying arteries to enable concurrent hemodynamic measurements across multiple levels of the *neurovascular complex*. Here, we apply this method across several levels of the vascular hierarchy, from the basilar and posterior cerebral arteries to pial arteries, pial veins, and major draining venous sinuses such as the straight and superior sagittal sinuses.

Stimulus-locked velocity increases were observed in individual vessels within each subject. Arterial responses occurred rapidly and nearly simultaneously across upstream and pial arterial segments, whereas venous responses exhibited temporal delays relative to the arterial responses. The average arterial-to-venous response time delay differed between the onset and offset phases, measuring approximately 4.1 s during response onset and 1.3 s during response offset, and also varied across different vessel segments. A simplified Windkessel model—used to interpret these findings—predicted similar inflow–outflow delays, suggesting that these temporal differences may reflect the transient filling and draining dynamics within compliant vascular compartments intervening between supplying arteries and draining veins. Our results demonstrate the feasibility of using dynamic PC-fMRA to measure vessel-specific hemodynamic responses across the human neurovascular complex. By providing quantitative measurements of macrovascular responses that serve as inputs to and outputs from intervening vascular compartments, this approach offers a framework for probing the hemodynamics within these compartments and studying vascular network behavior during neuronal activation, which may also help improve the physiological interpretation of fMRI signals in humans.

## 1 Introduction

Increased neuronal activity in the brain is accompanied by changes in cerebral blood flow (CBF) to ensure that an adequate supply of oxygen and nutrients is delivered and waste is cleared to maintain normal brain function (Iadecola, 2017; Willie et al., 2014). This active vascular response to neuronal activity causes an increase in blood delivery—functional hyperemia—is the most salient physiological manifestation of neurovascular coupling (NVC). NVC refers to mechanisms of communication between neurons and parenchymal microvessels that regulate local vascular tone and blood flow, and this tight cooperation motivated the concept of the neurovascular unit (NVU) (Iadecola, 2017). Recent findings, however, suggest that the NVU operates within a larger, interconnected vascular hierarchy: microvascular responses are embedded within a cerebrovascular tree whose macrovascular segments can influence and reflect neural activity at spatial scales far beyond the local microvasculature. This broader viewpoint is captured by the concept of the *neurovascular complex* (Schaeffer & Iadecola, 2021), which emphasizes that coordinated vascular responses can extend across multiple scales and levels of the vascular network: upstream from the local microvasculature to pial arteries and further to major intracranial and even extracranial arteries, and downstream to veins and sinuses. From this perspective, hemodynamics during neuronal activation cannot be fully understood as the response of microvessels alone—it rather emerges from coordinated changes within the larger interconnected network that redistributes blood flow and blood volume across multiple vascular compartments (Ji et al., 2026). Characterizing these dynamics at the network level is therefore critical for understanding human brain hemodynamics.

*In-vivo* microscopy in animal models has provided fundamental insights into NVC and how the microvascular network influences the hemodynamic responses (Ji et al., 2026). Studies have shown that neuronal activation-induced vasodilation is observed not only in microvessels near the activated site, but also in the upstream feeding arteries through retrograde propagation via signaling along the endothelial cells (Chen et al., 2011; Iadecola, 2017; Iadecola et al., 1997; Schaeffer & Iadecola, 2021). However, the limited field of view (FOV) of microscopy restricts its ability to characterize the full spatial extent of propagation and the integrated hemodynamics from distant arterial and venous segments throughout large-scale brain vascular networks. Consequently, our understanding of neurovascular responses integrated across a vascular network as large as that of the human brain is limited.

The unknowns matter, because local hemodynamic responses in the microvasculature are inherently shaped by the macrovasculature—pressures and impedances in feeding arteries and draining veins—which can strongly influence blood flow and volume changes throughout the network (Lorthois et al., 2011; Pfannmoeller et al., 2021). These knowledge gaps also have implications for the interpretation of functional MRI (fMRI), which remains the most widely used approach for studying human brain activity noninvasively (Attwell & Iadecola, 2002; Bandettini et al., 1992; Kwong et al., 1992; Ogawa et al., 1992). All fMRI methods (Belliveau et al., 1991; Kwong et al., 1992; Lu et al., 2003; Williams et al., 1992) in use today track brain activity through the associated hemodynamic changes in blood flow, blood volume, and blood oxygenation, thus a comprehensive understanding of the vascular responses underlying these changes is essential for accurate interpretation of the fMRI signals. For example, vascular anatomic network (VAN) models and related frameworks combine realistic vascular geometry and dynamics to simulate and, thereby, interpret fMRI signals (Boas et al., 2008; Gagnon et al., 2015; Gagnon et al., 2016; Hartung et al., 2026; Pfannmoeller et al., 2021). However, modern VAN models typically represent a localized microvascular network—including arterioles, capillaries, and venules—within a single brain location, i.e., one fMRI voxel, and do not explicitly include the macrovascular networks that supply and drain the microvascular networks. Without detailed measurements regarding the upstream and downstream boundary conditions that constrain local pressure gradients and flow distribution, these values are often assumed fixed (Gagnon et al., 2015; Hartung et al., 2022; Hartung et al., 2026). In addition, many model parameters derived from invasive animal measurements—such as vascular geometry (e.g., vessel density) and topology (e.g., the arteriole-to-venule ratio)—may not generalize to human vascular physiology (Hartung et al., 2022; Hartung et al., 2026). Directly measuring human hemodynamics at multiple vascular stages—rather than focusing exclusively on fMRI signals that reflect changes within the local microvasculature—therefore can provide more accurate inputs to these biophysical models as well as a means to directly test model predictions as part of the model validation. A range of measurement modalities has been used to probe human vascular responses. Functional transcranial Doppler (fTCD) (Aaslid, 1987) provides high temporal resolution measurements of blood velocity in a small set of large intracranial arteries accessible to ultrasound, including first- or second-order branches of the anterior cerebral artery (ACA), middle cerebral artery (MCA), and posterior cerebral artery (PCA) (Aaslid, 1987; Conrad & Klingelhofer, 1989; Droste et al., 1989; Kelley et al., 1992; Phillips et al., 2016; Smith et al., 2008; Sturzenegger et al., 1996). The high temporal resolution and non-invasive nature of fTCD make it a useful tool for assessing cerebrovascular function and how neuronal activity and local hemodynamic responses generate global changes in blood flow measured in far upstream arteries. However, its spatial resolution limits this technique to measurements of the major cerebral arteries, and the accuracy of the fTCD measurement is dependent on the training and experience of the technicians (Pan et al., 2022). MR angiography (MRA) techniques can provide much higher spatial resolution compared with fTCD, as well as the ability to target a larger set of intracranial vessels, and have been mainly used for measuring anatomy or baseline physiology of vessels in clinical settings (Hibert et al., 2021; Rivera-Rivera et al., 2020). A new set of imaging tools known as “functional MRA” (fMRA) (Park et al., 2018) has adapted MRA techniques to measure dynamic vascular responses to neural activity, including functional Time-of-Flight (fTOF) (Belle et al., 1995; Bizeau et al., 2018; Cho et al., 2008; Cho et al., 2012) and phase-contrast functional MRA (PC-fMRA) (Bizeau et al., 2018; Chen et al., 2021; Hu et al., 2025; Kang et al., 2010; Park et al., 2018). The fTOF technique utilizes the velocity-related inflow enhancement effect (Wehrli, 1990) to measure the intensity changes induced by blood velocity increases initiated by neuronal activity. Although fTOF exhibits high sensitivity to velocity changes, it is mainly used as a qualitative method, and the measures it provides can be biased by partial-volume effects (Bizeau et al., 2018). Thus, this approach trades off specificity for sensitivity. The PC-fMRA technique leverages the conventional phase-contrast MRA (PC-MRA) method to directly quantify blood velocity changes induced by neuronal activation. In PC-MRA, a pair of bipolar velocity-encoding gradients encodes blood velocity into the phase of the MR signal, with the resulting phase shift being proportional to the linear velocity of moving spins; this phase shift can then be converted into quantitative velocity values (Moran, 1982).

Recently, a dynamic “vessel-specific” (Chen et al., 2021) PC-fMRA method was introduced in the context of preclinical MRI in mice to enable quantitative velocity measurements at temporal resolutions comparable to conventional block-design fMRI studies while targeting specific vessels and has been extended to human brain using clinical MRI scanners (Hu et al., 2025). A conceptual outcome of this work in the human brain is that stimulus-evoked velocity responses measured in a large feeding artery segment—located centimeters upstream of the activated cortex—can plausibly reflect passive *network-level* consequences of downstream resistance changes rather than local active dilation at the measurement site. In this view, local arteriolar and microvascular dilation within activated tissue increases perfusion through decreased parenchymal blood-flow resistance, and mass balance within the interconnected vascular tree necessitates corresponding flow changes in upstream feeding arteries and potentially in other connected segments. Such observations of *network-level effects* imply that hemodynamic changes can be spatially interdependent across multiple vascular segments, and that interpreting hemodynamic signals benefits from a framework that explicitly accounts for the coupling of hemodynamics within multiple levels of the full vascular network.

While a response in large upstream arteries, such as the branches of the PCA reported by Hu et al. (2025), may reflect velocity changes as a mass-balance consequence of downstream resistance changes, similar blood velocity dynamics should not be assumed to occur across all vascular segments. Classical Windkessel models (Mandeville et al., 1999; Westerhof et al., 2009) describe the vascular system as a resistor-capacitor network, in which changes in vascular resistance alter pressure and flow distribution while vascular capacitance allows blood volume to change. The relationship between upstream arterial inflow and downstream venous outflow is thus shaped by the compliant microvascular compartments between them. This principle is also central to the “Balloon model” of the fMRI response, which emphasizes the dynamically-changing relationship between arterial inflow, blood volume, and venous outflow including transient decoupling and recoupling between these hemodynamic components (Buxton et al., 1998; Mandeville et al., 1999; Polimeni & Lewis, 2021). Concurrent measurements across both arterial and venous compartments are therefore needed to understand how stimulus-evoked hemodynamic changes temporally evolve across the vascular network during neuronal activation to properly interpret the fMRI response in all its complexity.

Motivated by this need, this study aims to use the dynamic PC-fMRA (Hu et al., 2025) technique to concurrently measure the hemodynamic response to neuronal activation at multiple stages of the human neurovascular complex—including major supplying arteries, pial arteries, pial veins, and major draining veins and sinuses. Specifically, we use visual stimulation to investigate the vessels that supply and drain the human visual cortex. By measuring the coupled dynamics across arterial and venous pathways, we aim to provide a preliminary *network-level* characterization of vascular responses that can further inform understanding of coordinated response across the vascular network in the human brain that occurs with neuronal activation.

## 2 Methods

### 2.1 Study participants

Five healthy adults volunteered to participate in the study. All volunteers provided written informed consent prior to scanning, in accordance with our approved protocol and the policies of our institution’s Human Subjects Research Committee. The experiments were conducted on a whole-body 7T scanner (Terra, Siemens Healthineers, Erlangen, Germany) using an in-house-built 64-channel brain receive coil array and “birdcage” volume transmit head coil (Mareyam et al., 2020).

### 2.2 MR imaging data acquisition

To enable concurrent measurements from multiple vessels, a single oblique-axial slice that intersected several vessels of interest was required. Anatomical images, including multi-echo MPRAGE and 3D TOF-MRA (Figs. 1A and 1C), were first acquired to aid in prescription of the PC-fMRA acquisition. An oblique-axial slice was then positioned to intersect the basilar artery (BA), straight sinus (STS), superior sagittal sinus (SSS), and pial vessels overlying the occipital cortex, including pial arteries (PA) and pial veins (PV) (Figs. 1A and 1B). To compare with our previous measurements, we also acquired a second oblique-coronal slice prescribed approximately perpendicular to the bilateral second-order branch (P2) segments of the PCA (Figs. 1C and 1D). The PC-fMRA single-slice acquisitions used standard one-sided velocity encoding with a single velocity encoding direction (the through-plane direction) and one velocity encoding (VENC) value (Moran, 1982) with the following parameter values: FOV = 180 × 180 × 2 mm^3^, in-plane resolution = 0.8 × 0.8 mm^2^, slice thickness = 2 mm, VENC = 60 cm/s, flip angle = 13°, GRAPPA acceleration factor = 2, TR/TE = 15.70/4.62 ms. The pulse sequence alternated between acquiring one k-space line for an image without velocity encoding and one k-space line for an image with velocity encoding, as in conventional PC-MRA. Effective temporal resolution corresponded to the time required for acquiring the image pair, i.e. 2.01 s. Example phase-difference images— which can then be converted into velocity maps—from the two PC-fMRA slices are presented in Figs. 1B and 1D, and the positions of different vessels are indicated by white arrows. For each of the two slice prescriptions, three PC-fMRA runs were acquired. In two of the five subjects, three additional lower-VENC PC-fMRA runs were acquired with VENC = 10 cm/s using the slice prescription targeting the PAs and PVs, to improve sensitivity to the relatively low velocities in these vessels. The PC-fMRA acquisitions were not cardiac-gated, and since the achieved temporal resolutions were approximately two cardiac cycles under normal conditions (assuming a typical cardiac cycle frequency of about 1 Hz), the measured velocities represent averages across the cardiac cycle. A high-resolution eight-echo FLASH (TR/TE1/ΔTE = 62/4.32/6.44 ms, in-plane resolution = 0.14 × 0.14 mm^2^, slice thickness = 2 mm, flip angle = 60°) was also acquired in the same slice prescriptions to distinguish between pial arteries and veins based on their distinct T_2_* values (Fig. 1E).

**Figure 1:**
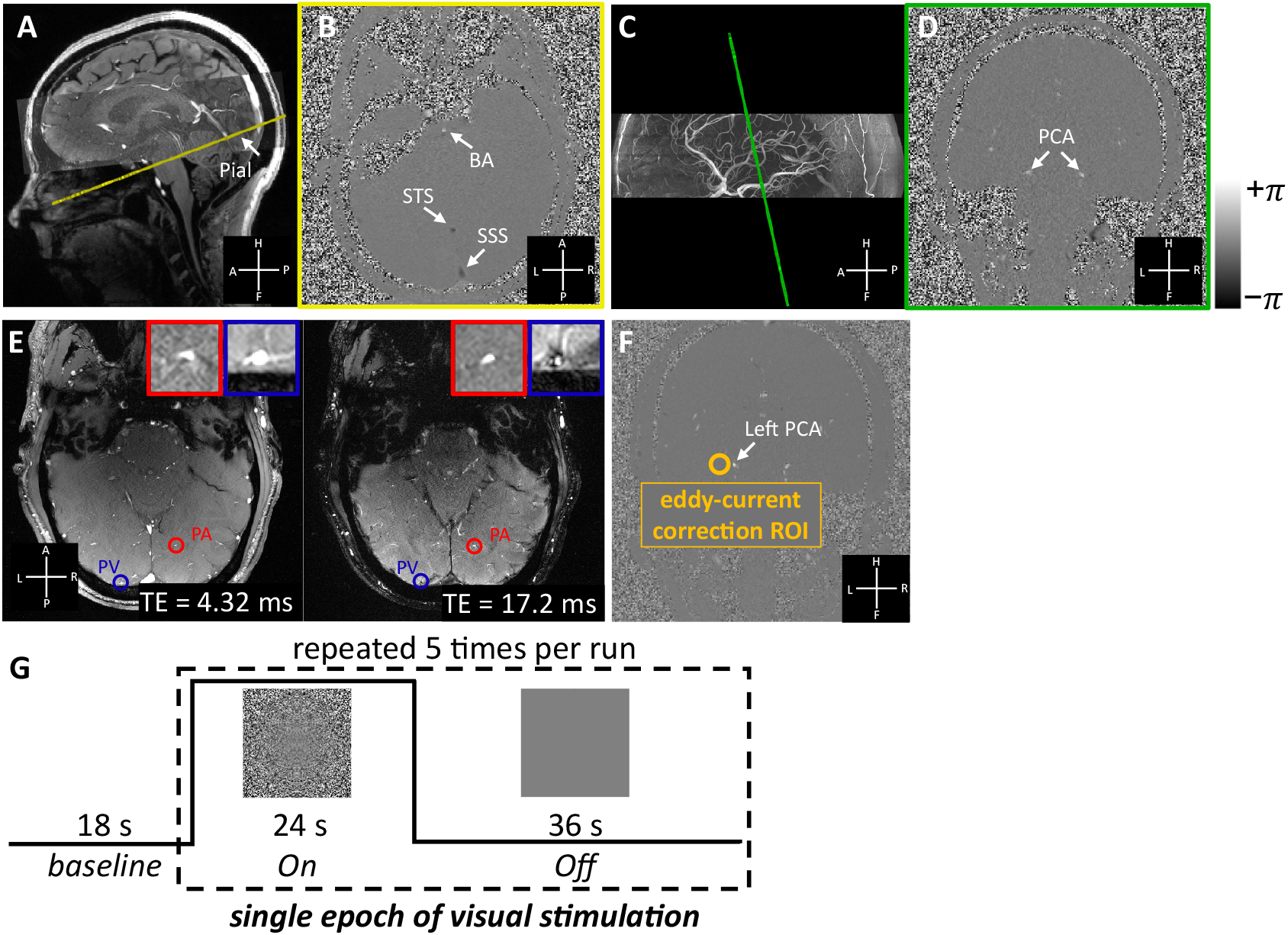
Experimental design and prescription of the PC-fMRA acquisition. Positions of the two slices (indicated by yellow and green, respectively) are shown on either multi-echo MPRAGE or 3D TOF-MRA data (A, C). The first, oblique-axial slice (A, B) was prescribed to intersect multiple vessels for concurrent velocity measurement including the basilar artery (BA), straight sinus (STS), superior sagittal sinus (SSS), and pial vessels (Pial)—both pial arteries (PA) and pial veins (PV)—within the occipital cortex. The second, oblique-coronal slice (C, D) was prescribed as perpendicular to the “P2” segments of posterior cerebral arteries (PCA). The positions of individual vessels are indicated by white arrows. Images at different TEs from a multi-echo FLASH were used to distinguish between pial arteries and veins based on differences in T_2_* decay of arterial and venous blood. Note that the images acquired at different echo times are displayed with different color scales for better visualization. Representative PA and PV examples are shown as insets with red and blue borders, respectively (E). The PA shows higher intensity than the surrounding tissue at both echo times, whereas the PV shows higher intensity than the surrounding tissue at the shorter echo time but lower intensity at the longer echo time. Velocities were corrected for biases induced by eddy currents using background phase information from neighboring ROIs: an example ROI used for the correction of the left PCA is shown in (F) (orange circle). The legends in panels (A)–(F) indicate the anatomical axes (A: anterior, P: posterior, H: head, F: foot, L: left, R: right). The stimulation paradigm consisted of an initial 18-s baseline period followed by five epochs of counterphase-flickering scaled-noise stimuli per run (a 24-s ON period and a 36-s OFF period) (G).

### 2.3 Eddy current correction

The relatively large gradient strengths and slew rates used for velocity encoding induced eddy currents that manifested as VENC-dependent phase offsets in static tissue in the phase-difference images. These unwanted phase effects could bias flow velocity measures (Pelc et al., 1994). Thus, eddy-current biases were estimated and corrected by extracting the background phase from a region of interest (ROI) in tissue devoid of visible vessels adjacent to each targeted vessel (Fig. 1F), similar to a previously described correction method (Hibert et al., 2021; Pelc et al., 1994).

### 2.4 Experimental design

The stimulation protocol consisted of a standard block-design paradigm beginning with an 18-s baseline period followed by five epochs of 24-s ON periods in which 8-Hz counterphase-flickering logarithmically-scaled spatial-noise stimuli were presented separated by 36-s OFF periods in which a neutral gray screen was presented (Fig. 1G). A central red dot was displayed throughout the stimulation as a fixation target; participants were instructed to maintain fixation and press any button on an MRI-safe button box when the dot randomly turned either brighter or darker. The visual stimulus was implemented with `Psychtoolbox-3` (https://www.psychtoolbox.net) (Kleiner et al., 2007) in MATLAB (The MathWorks, Natick, MA). The total scan time per run was 5 min 18 s.

### 2.5 MR imaging data analysis

In-plane motion was estimated on the magnitude-valued time-series data from the PC-fMRA scan using `3dAllineate` from AFNI (https://afni.nimh.nih.gov/) (Cox, 2012), with the frame acquired in the middle of the three runs as the reference to correct both within-run and across-run motion. The estimated transformation matrices were then applied to the corresponding phase-difference maps, which were then converted into velocity maps and averaged across the three runs. These averaged time-series data were analyzed using a standard General Linear Model (GLM) with a canonical Gamma-variate hemodynamic response function (HRF)—which is intended to model the BOLD response but here is used to model the blood velocity response—implemented in FSL `FEAT` (Woolrich et al., 2001) to generate Z-score maps, which were then thresholded using clusters determined by *z* > 3.1 and a cluster significance threshold of *p* < 0.05 (Worsley, 2001) to produce the activation maps. Although a small number of PC-fMRA voxels intersecting the targeted blood vessels were above the set activation threshold, this correction for multiple comparisons was helpful in automatically removing isolated false-positive activations seen in tissue regions that did not contain vessels. The GLM analysis was mainly used to automatically identify responsive voxels, after which the time-series data were further evaluated. After eddy-current correction, the time-series of velocities from the voxel with the maximum velocity in each of the vessels were extracted and interpolated to a temporal sampling grid of 0.1s spacing using linear interpolation. Trial-averaged responses were then computed.

To quantify response timing, the 50% transition time (*t_50_*) was estimated from the trial-averaged velocity response of each individual vessel. The onset and offset transitions were analyzed separately and fitted with a four-parameter logistic sigmoid using nonlinear least-squares optimization. For the onset transition, the model was 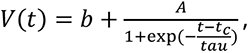, whereas the sign of the exponential term was reversed for the offset transition. Here, *b* represents the baseline, *A* the response amplitude, *t_c_* the midpoint of the fitted transition, and *tau* its temporal width. The 50% transition time was defined as *t_50_* = *t_c_*, corresponding to the time at which the fitted response reached the midpoint of its fitted response amplitude. For estimation of onset and offset timing, sigmoid fitting was performed over 0–18 s after stimulus onset and from 5 s before to 18 s after stimulus offset, respectively. This procedure provided vessel-specific *t_50_* estimates for subsequent comparison of arterial and venous response timing.

## 3 Results

Figure 2 shows the vessel-specific visual-stimulus-evoked blood velocity responses from three example subjects. Across all subjects, velocity increases in the BAs, PCAs, PAs and PVs were observed in the trial-averaged responses. These findings were further confirmed by GLM analysis of the time-series data (thresholded using clusters determined by *z* > 3.1 and a cluster significance threshold of *p* < 0.05), which consistently detected stimulus-locked velocity responses in these individual vessels. Peak stimulus-evoked velocity changes, calculated relative to the velocity at stimulus onset (*t* = 0) using the peak velocity of the trial-averaged response associated with the visual stimulus, were approximately 5– 15% in the BA and PCA. Larger relative changes were observed in the smaller pial vessels, with increases of approximately 50–100% in the PA and PV. In Subject 1 and Subject 2, velocity increases were observed in both the STS and SSS, whereas in Subject 3, it was more pronounced in the STS than in the SSS, possibly reflecting intersubject variability in the cortical regions drained by these sinuses.

**Figure 2:**
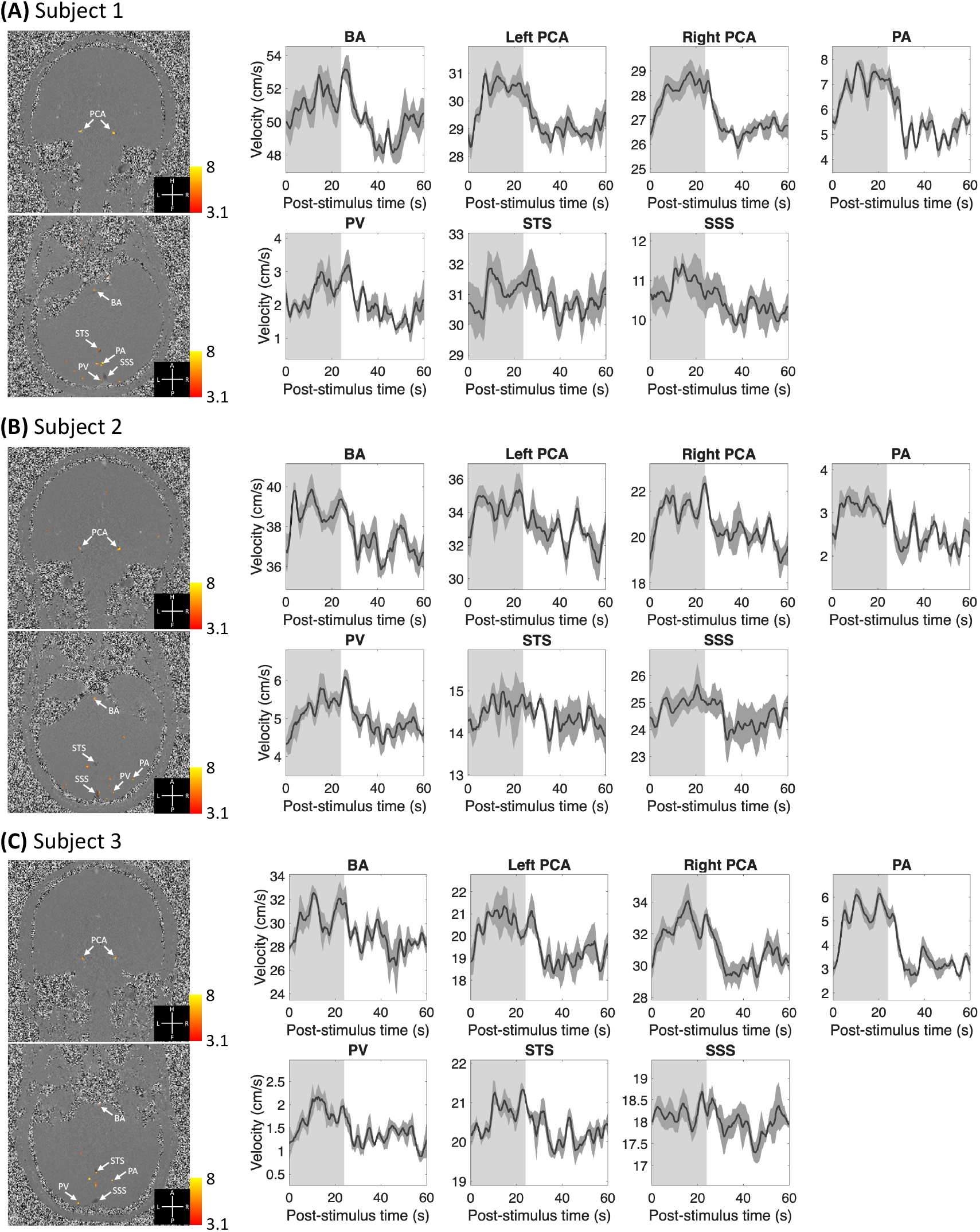
Vessel-specific blood velocity responses measured in three example subjects (healthy volunteers). Z-score maps (thresholded using clusters determined by *z* > 3.1 and a cluster significance threshold of *p* < 0.05), which indicate the activated regions, were overlaid on example single-frame phase-difference images (first column). Trial-averaged responses (shading represents standard errors calculated across all 15 trials) from the voxel with the maximum velocity from each individual vessel are shown on the right. The light gray and white regions in the plots represent the ON and OFF periods, respectively. (A) Subject 1. (B) Subject 2. (C) Subject 3.

Relative latencies in the venous responses were also observed in these subjects (Fig. 2). The response onsets appeared similar across the imaged arterial vessels (BA, PCA, and PA), whereas the venous responses generally appeared later, including those in the draining veins and sinuses. We further characterized the arterial–venous latency in Figure 3. Trial-averaged velocity responses from individual PAs (Fig. 3A) and PVs (Fig. 3B) showed systematically later transitions in the PVs. The 50% transition time *t_50_* was estimated for each vessel. Across the measured vessels (*N* = 10 PAs and *N* = 10 PVs from all 5 subjects), PVs exhibited significantly later *t_50_* values than PAs during both onset (Fig. 3C; two-sided Welch’s two-sample *t*-test, *p* < 0.001) and offset (*p* < 0.05). Onset transition times were more consistent across PAs, whereas PVs exhibited a broader distribution, indicating greater inter-vessel variability in venous response timing. The venous latency was also evident in the group-averaged normalized responses (Fig. 3D), with PA–PV temporal delays of approximately 4.1 s during response onset and 1.3 s during response offset. The difference between the onset and offset delays suggests some asymmetry in the relationship between arterial and venous dynamics during the two transient phases of the hemodynamic response.

**Figure 3:**
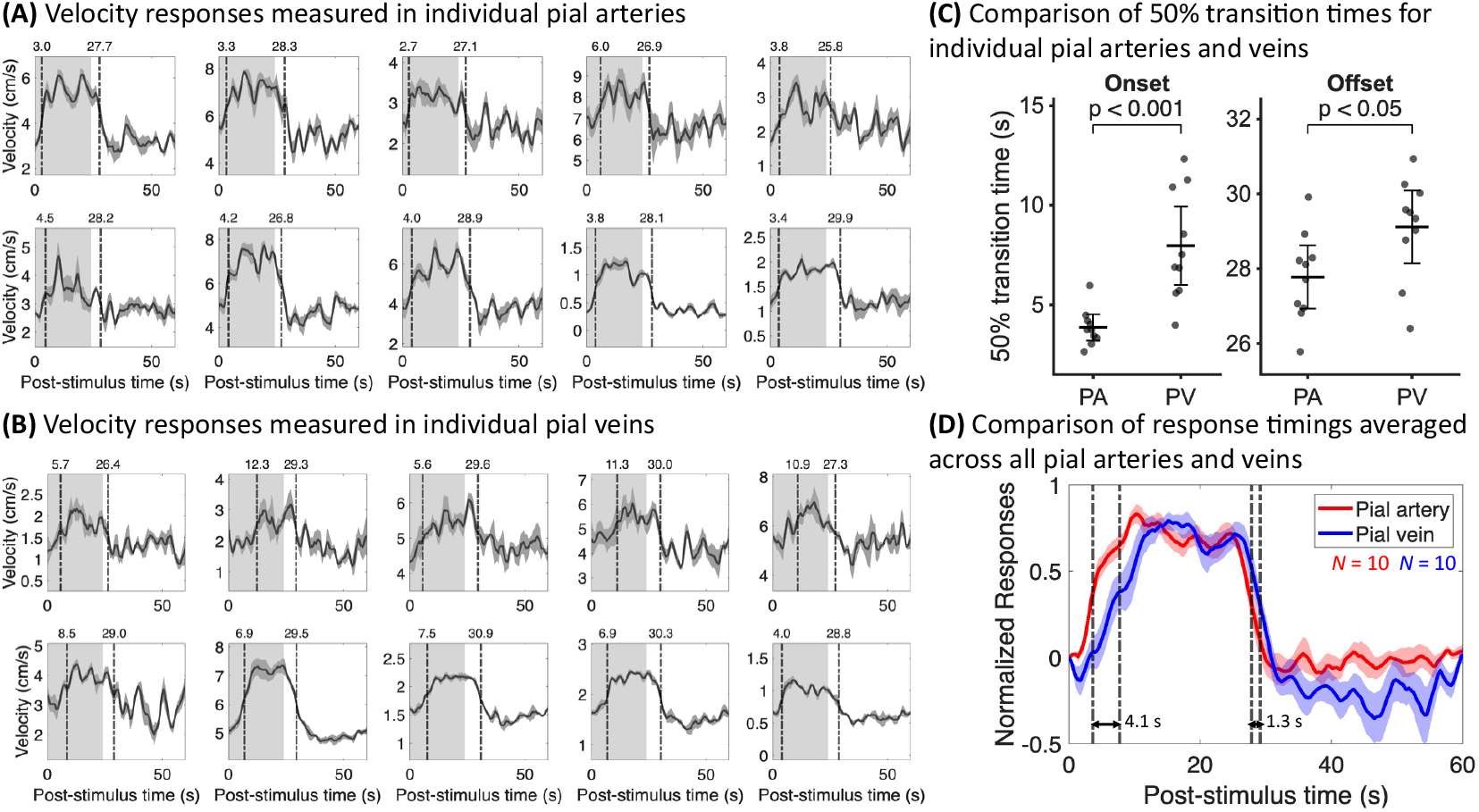
Temporal delay between arterial and venous responses. Trial-averaged velocity responses in individual pial arteries (A) and veins (B) from all 5 subjects (shading represents standard errors calculated across all 15 trials). The last four PAs and PVs were measured with VENC = 10 cm/s, yielding higher SNR. The light gray and white regions in the plots represent the ON and OFF periods, respectively. For each vessel, the vertical dashed line indicates the 50% transition times (*t_50_*) for the onset and offset periods, with the corresponding *t_50_* value shown above each plot. (C) Comparison of individual onset and offset *t_50_* values between PAs and PVs. Individual vessel measurements are shown as points, with black horizontal lines indicating the group means and error bars indicating 95% confidence intervals. PVs exhibited significantly later transition times than PAs during both onset (two-sided Welch’s two-sample *t*-test, *p* < 0.001) and offset (*p* < 0.05). (D) Group-averaged normalized responses across all measured PAs and PVs (*N* = 10 PAs and *N* = 10 PVs from all 5 subjects; shading indicates standard errors across vessels). The vertical dashed lines indicate the 50% transition times of the group-averaged PA and PV responses. The corresponding temporal delays were approximately 4.1 s during response onset and 1.3 s during response offset, as indicated by the horizontal black arrows.

## 4 Discussion

In this study, we extended our vessel-specific PC-fMRA approach (Hu et al., 2025) to quantitatively measure and compare the timing of blood velocity responses induced by visual stimulus across multiple stages of the neurovascular complex, including major supplying arteries, pial arteries, pial veins, and major draining veins and sinuses. Consistent velocity increases were measured in individual vessels within individual subjects, demonstrating the feasibility of this approach for comparing vascular responses at multiple targeted locations across the neurovascular complex. These measurements extend the study of neurovascular coupling beyond local tissue responses and suggest that the measured macrovascular dynamics may provide meaningful information about how blood delivery and drainage dynamics emerge from the interactions across the vascular network during neuronal activation.

In particular, we observed rapid velocity increases in upstream supplying and pial arteries, together with a later response pattern in downstream pial veins and draining sinuses. It is known that neuronal activity-induced vasodilation first occurs in local parenchymal microvessels and arterioles then propagates upstream to larger arteries through signaling along endothelial cells (Chen et al., 2011; Chen et al., 2014; Iadecola et al., 1997). Thus, the observed blood velocity increases in the PAs near the activated tissue may reflect active vessel dilation triggered via this well-known mechanism of neurovascular coupling and active vascular coordination, though propagation of dilation to vessels of this caliber remains to be demonstrated. Furthermore, the responses in further upstream arteries, such as the BAs and PCAs, were also relatively fast, with no obvious differences in response timing relative to the PA responses (Fig. 2). Given the reported speed of retrograde propagation of vasodilation signaling along surface arterioles (∼0.24 cm/s) (Chen et al., 2011; Hillman et al., 2007; Tian et al., 2010), it is unlikely that these upstream responses were driven primarily by this signaling pathway. (Retrograde propagation from activated brain tissue closer to the upstream arterial segments we measured—including subcortical regions such as the lateral geniculate nucleus (LGN) (Chen et al., 1998), which are also activated by visual stimulation—remains a possibility and may contribute some of this response, however the LGN tissue volume and thus blood volume is far less than that of visual cortex.)

Our observation is consistent with our hypothesis of a *network-level* effect (Hu et al., 2025): the observed velocity increases in the far-upstream arteries may be passive in nature. In this context, “passive” refers to an upstream velocity increase that arises as a hemodynamic consequence of increased downstream blood flow, rather than from NVC-mediated vasodilation in these arteries. Instead, local arteriolar and microvascular dilation within activated tissue decreases its vascular resistance, thereby increasing local blood flow in the tissue. Through conservation of mass within the interconnected vascular tree, local increased blood flow in the tissue would necessarily lead to increased blood flow in upstream branches of the supplying arteries. If these supplying arteries do not dilate, then this increased blood flow (i.e., volumetric flow) would be equivalent to increased blood linear velocity, as measured here. Autoregulatory responses may nevertheless occur concurrently in the upstream vasculature. Cerebral autoregulation refers to vascular mechanisms that adjust cerebrovascular tone in response to changes in perfusion pressure (Faraci & Heistad, 1990; Panerai, 1998; Roy & Sherrington, 1890; Strandgaard & Paulson, 1984; van Beek et al., 2008; Willie et al., 2014). Such upstream vascular responses could be triggered by changes in perfusion pressure associated with the reduction in vascular resistance in the occipital cortex during neuronal activation, or by flow-mediated dilation. However, autoregulatory responses typically engage with a latency of several seconds (Claassen et al., 2021). The rapid onset of the velocity increases observed in our data therefore suggests that autoregulation is unlikely to be the primary mechanism underlying the earliest component of the observed responses, although it may modulate the response during later phases (Hu et al., 2025).

Latencies of the venous responses relative to the arterial responses were also observed in the measured data (Figs. 2 and 3). One potential framework for interpreting this behavior is the classic Windkessel model (Mandeville et al., 1999; Suarez et al., 2021), as illustrated in Figure 4. Fig. 4A shows a schematic illustration of a three-element Windkessel model. Blood flow from the supplying arteries runs through the intervening vascular network before draining into the veins. During sustained neuronal activation, pial arteries may dilate, leading to a time-varying decrease in arterial resistance, represented by *R*_a_(*t*). The intervening vasculature (i.e., any vascular segments between the measured artery and measured vein) may also dilate, resulting in increased blood volume; this process is modeled using a resistor–capacitor circuit, represented by fixed values for *R*_w_ and *C*_w_. The upstream boundary pressure is also fixed at *P_0_*. The time-varying pressure at the branch point is denoted by *p*(*t*). Based on mass conservation, the relationship between the inflow dynamics (*F*_IN_) and the corresponding outflow dynamics (*F*_OUT_) can be described

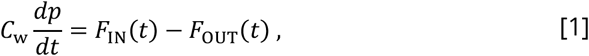

**Figure 4:**
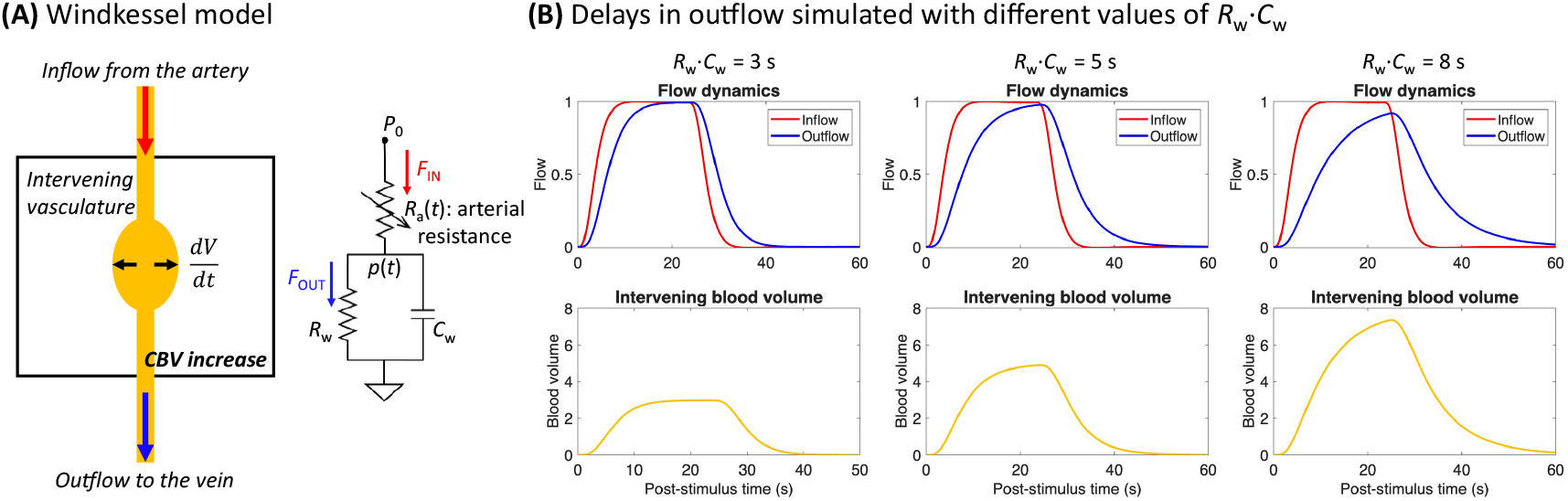
Simulation of a three-element Windkessel model. (A) Schematic illustration of the Windkessel model. Blood flow from the supplying arteries runs through the intervening vascular network before draining into the veins. During sustained neuronal activation, pial arteries may dilate, leading to a decrease in arterial resistance, represented by *R*_a_. The intervening vasculature may also dilate, resulting in increased blood volume; this process is modeled using a resistor–capacitor circuit, represented by *R*_w_ and *C*_w_. The upstream boundary pressure is fixed at *P_0_*. The time-varying pressure at the branching point is denoted by *p*(*t*). Given the inflow dynamics (*F*_IN_) driven by changes in *R*_a_, the corresponding outflow dynamics (*F*_OUT_) and blood volume changes can be simulated. Simulations of outflow delays with different *R*_w_⋅*C*_w_ values (3, 5 and 8 s), along with the corresponding blood volume changes, are also shown (B).

where, by the fluidic version of Ohm’s law, 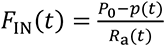 and 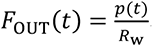, which leads to

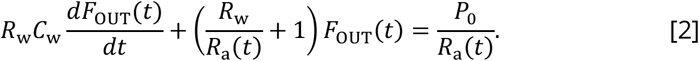

Given the inflow dynamics *F*_IN_(*t*), which are driven by changes in *R*_a_, the corresponding outflow dynamics and blood volume changes in the intervening vasculature can be simulated numerically. For fixed values of *R*_w_ and *C*_w_, because the delay between inflow and outflow is related to the product of *R*_w_ and *C*_w_ in the Windkessel model (Mandeville et al., 1999), a fixed temporal delay between inflow and outflow is observed resulting in a symmetry in the relationship between arterial and venous dynamics during these two transient phases of the hemodynamic response (Fig. 4B). The temporal ordering of arterial and venous responses is consistent with the dynamics of the blood passing through the vascular network, and supports the idea that arterial inflow and venous outflow are linked through an intervening vascular compartment whose volume changes during activation. Intuitively, the temporal delay between arterial and venous hemodynamic response timing represents the time taken by blood filling the slowly-expanding “balloon” that connects the measured input and output vessels, consistent with mass conservation.

When comparing the responses in PAs and PVs, different temporal delays were observed during the onset and offset periods of the visual stimulus (Fig. 3C). To further characterize these delays, we used the normalized response in PAs as the input to the Windkessel model and identified the output that best matched the normalized responses in PVs based on temporal cross-correlation. This analysis provided estimates of the corresponding *R*_w_⋅*C*_w_ values of the Windkessel model. By maximizing the correlation separately for the onset and offset periods, we obtained *R*_w_⋅*C*_w_ values of 4.1 s and 1.1 s, respectively (Fig. 5A), in good agreement with the directly estimated temporal delays of 4.1 s and 1.3 s. To examine how response timing varied along the measured vascular hierarchy, we next used the PCA response as a common upstream reference and estimated the temporal delays of downstream PA, PV, and venous sinus responses relative to it using data from the onset periods. These delays were 1.1 s, 5.3 s and 7.6 s respectively (Fig. 5B). The relatively small PCA-to-PA delay may partly reflect the fact that the measured PAs are not necessarily the earliest responding arterial segments supplying the activated regions. The progressively larger delays observed in the PVs and venous sinuses are consistent with additional temporal filtering within the intervening vascular network, as described by the Windkessel framework. From Equation [2], the temporal properties of the outflow relative to the inflow depend on the values of *R*_w_, *C*_w_, and *R*_a_(*t*) . Thus, differences in the downstream venous delays may reflect regional differences in the properties of the intervening vascular compartments, whereas the different onset and offset delays may reflect temporal changes in the resistance and compliance dynamics of the vascular system. The asymmetry in the temporal delay during onset and offset periods suggests that the vascular system may operate under different dynamic conditions during the rising and recovery phases of the response. This observation is notable because onset–offset asymmetries have long been recognized in conventional hemodynamic signals, including BOLD fMRI—classically modeled as differences in “inflation” and “deflation” time constants in the context of the Balloon model (Buxton et al., 2004)—but their physiological basis remains incompletely understood.

**Figure 5:**
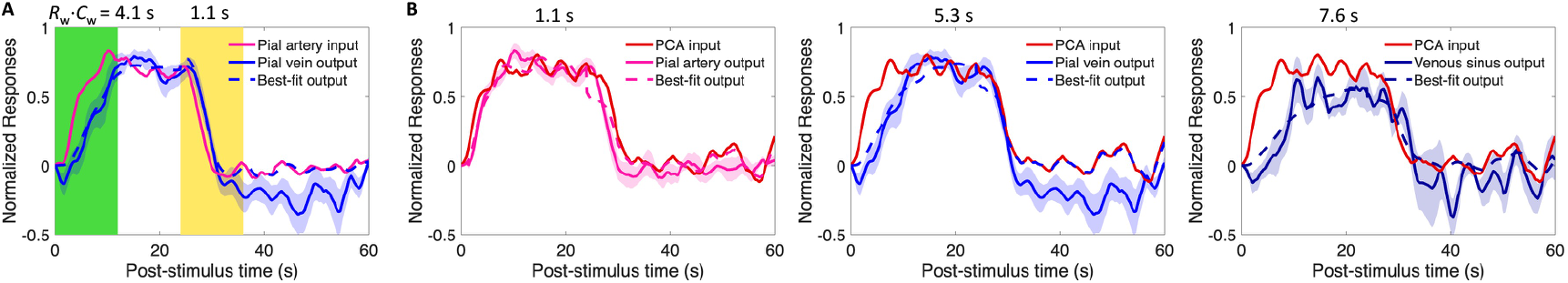
Temporally and spatially varying delays. (A) Using the normalized pial arterial response as the input, the best-fit Windkessel model whose output most closely matched the normalized pial venous responses was identified, thereby providing estimates of the corresponding *R*_w_⋅*C*_w_ values. These values were 4.1 s and 1.1 s for the onset and offset periods, respectively, in good agreement with the directly estimated temporal delays. (B) Using the measured PCA response as a common upstream reference, distinct temporal delays during the onset period were estimated for downstream vessels: 1.1 s for pial arteries, 5.3 s for pial veins, and 7.6 s for venous sinuses. Responses were averaged across measured vessel segments from 5 subjects, including 10 bilateral PCA segments, 10 pial arterial segments, 10 pial venous segments, and 7 venous sinus segments from the STS and SSS.

By directly measuring arterial input and venous output in individual vessels, phase-contrast fMRA may provide a means to quantify vascular network-level and microvascular dynamics within the activated tissue supplied and drained by the measured arteries and veins, respectively. Specifically, concurrent measurements of arterial inflow and venous outflow may provide a basis for estimating an effective “transfer function” of the intervening microvascular compartments—that is, how stimulus-evoked inflow changes are transformed into downstream venous responses through the tissue. This *input–output* perspective could potentially help interpret hemodynamic phenomena that are difficult to access using conventional fMRI approaches. Similar concepts have been explored previously by using CBF estimates provided by arterial spin labeling (ASL) as inputs to Balloon models (Obata et al., 2004) to understand the post-stimulus undershoot in BOLD signals (Chen & Pike, 2009). In a related but distinct approach, dynamic susceptibility contrast MRI (DSC-MRI) uses an arterial input function to characterize the contrast-agent bolus that is delivered to tissue and uses this to estimate perfusion through deconvolution (Ostergaard, Sorensen, et al., 1996; Ostergaard, Weisskoff, et al., 1996; Rosen et al., 1991). However, ASL is an indirect tissue-level perfusion measurement with its own signal-modeling complexities, and the DSC arterial input function is based on exogenous contrast-agent passage through the tissue, whereas here we are using blood velocity time-courses. In this sense, vessel-specific PC-fMRA may offer a complementary framework by directly measuring arterial inflow and venous outflow—albeit in individual vessels— thereby providing more specific information on how the intervening vascular network shapes the temporal dynamics of the measured hemodynamic responses. Such measurements may ultimately help infer tissue-level or microvascular hemodynamics from vessel-level or macrovascular hemodynamics—providing a bridge between vessel-specific and conventional fMRI signals by linking the vascular input, intervening network dynamics, and downstream venous output within a common physiological framework.

Several limitations should, however, be noted. The present interpretation relies on a simplified Windkessel model that does not capture the full structural and functional complexity of the cerebral vascular network, including heterogeneous vessel properties, distributed blood supply and drainage pathways and collaterals/shunts, and potential nonlinearities. For example, the measured arteries may not necessarily supply the same vascular territory drained by the measured veins, which limits inference about temporal properties between specific vessel pairs that we have measured. In addition, the measured pial arteries may not represent the earliest responding arterial segments supplying the activated regions, which limits the anatomical interpretation of the approximately 1.1-s temporal delay observed between the PCA and pial arterial responses (Fig. 5B). Furthermore, the present analysis uses only the temporal characteristics of the normalized blood-velocity responses. This approach is reasonable only if the actual volumetric-flow and velocity responses share similar temporal properties. However, in vessels where dilation may occur, such as the measured PAs and PVs, potential decoupling between volumetric flow and blood velocity (Proulx et al., 2025) could introduce bias into our estimates. Future work should combine vessel-specific volumetric-flow estimates—which can be made through joint estimation of linear velocity and vessel diameter changes (Varadarajan et al., 2025)—with more detailed vascular modeling, improved information regarding whether there is a direct anatomical pathway connecting the measured arterial and venous segments. By relating measured arterial-to-venous response time delays to the anatomical organization of the supplying and draining pathways including the vascular network topology and geometry, such efforts may clarify whether patterns in such temporal properties of hemodynamics across the network can serve to quantitatively characterize vascular network behavior and thereby identify hemodynamic interactions and other emergent behavior.

There are additional technical limitations to this study. Firstly, as a single-slice acquisition, motion correction via registration during post-processing is challenging since any through-slice motion cannot be corrected. During acquisition, three runs were acquired successively for each experiment within each individual experimental session, and the subjects were instructed to remain still during the session. Although this approach yielded good robustness for our experiments, combining our acquisition with prospective motion correction—either within-run or even across-runs—could potentially improve consistency and reduce variability (Balasubramanian et al., 2025). For example, the *AutoCorrect* framework (van der Kouwe et al., 2005; Wighton et al., 2024) can be employed to update slice prescriptions for each run to account for between-run head motion (Proulx et al., 2025). Secondly, a single slice was used to provide velocity measurements of multiple vessels, including the BA, PA, PV, STS, and SSS. The position and orientation of the slice relative to the trajectories of each vessel can affect the measured velocities, thus, the inter-subject variability in the prescriptions of slices may have a strong influence on the absolute values measured, making them not comparable between subjects. Also for this reason, even within a single slice, absolute velocity values may not be directly comparable across different vessels since each vessel likely intersects the slice with a different angle of obliquity. Thirdly, absolute velocities may also be underestimated due to partial-volume effects (Zong & Lin, 2019), particularly for the smaller PAs and PVs with a relatively large VENC. We minimized this issue by analyzing responses exclusively from the most intravascular voxel exhibiting the peak baseline velocity in each vessel. Finally, because our PC-fMRA acquisition consists of a single slice, we expect inflow enhancement or TOF effects, which will cause both the magnitude and phase of our data to be influenced by blood velocity and the distribution of blood velocities across the vessel lumen (e.g., laminar flow), resulting again in potential biases in the velocity quantification (Pelc et al., 1994; Proulx et al., 2026; Varadarajan et al., 2025). Further development of methods to fully account for these effects could potentially provide more precise and accurate velocity estimates from our measured data.

## 5 Conclusions

A dynamic vessel-specific PC-fMRA approach was used in this study to quantitatively measure blood velocity responses at multiple levels of the neurovascular complex in humans. Visual-stimulus-evoked responses were measured in individual arteries and veins within individual subjects. Arterial responses occurred rapidly, whereas venous responses were substantially delayed. Although further work is warranted to obtain more quantitative volumetric-flow measurements and a more complete reconstruction of the vascular network, the relatively long latencies of the venous responses suggest the presence of an intervening compliant vascular compartment between the measured arterial input and venous output. These results demonstrate that vessel-specific hemodynamic measurements across many levels of the vascular network may provide a useful framework for probing network-level effects on hemodynamics during neuronal activation and for improving the physiological interpretation of fMRI.

## Data and Code Availability

The de-identified data that support the findings of this study are available upon reasonable request from the corresponding author.

## Declaration of the use of AI

AI tools were used to check spelling and grammar.

## Author Contributions

Z.H.: Conceptualization, Methodology, Software, Formal Analysis, Investigation, Data Curation, Validation, Visualization, Writing – Original Draft & Review & Editing; G.A.H.: Methodology, Writing – Review & Editing; S.P.: Methodology, Writing – Review & Editing; D.V.: Methodology, Writing – Review & Editing; S.B.: Methodology, Writing – Review & Editing; B.R.: Methodology, Writing – Review & Editing; J.R.P.: Conceptualization, Methodology, Writing – Review & Editing, Supervision, Resources, and Funding acquisition.

## Acknowledgements

We would like to thank Estee Perelgut, Sarah Richter, Kyle Droppa and Julianna Gerold for their help with subject recruitment and MRI scanning support. This work was supported in part by the NIH NIBIB (grants P41-EB030006 and R01-EB019437), NCCIH (grant R01-AT011429), NINDS (grant R01-NS114526), by the *BRAIN Initiative* (NIH NINDS grants U19-NS123717 and U19-NS128613; NIH NIMH F32-MH125599), and by the MGH/HST Athinoula A. Martinos Center for Biomedical Imaging; and was made possible by the resources provided by NIH Shared Instrumentation Grant S10-OD023637.

